# Imaging toxin-induced neuroinflammation in mice using hyperpolarized ^13^C magnetic resonance spectroscopy

**DOI:** 10.1101/605568

**Authors:** Lydia M Le Page, Caroline Guglielmetti, Chloé Najac, Brice Tiret, Myriam M Chaumeil

## Abstract

Lipopolysaccharide (LPS) is a commonly used agent for induction of neuroinflammation in preclinical studies. Upon injection, LPS causes activation of microglia and astrocytes, whose metabolism alters to favor glycolysis. Assessing *in vivo* neuroinflammation and its modulation following therapy remains challenging, and new non-invasive methods allowing for longitudinal monitoring would be greatly valuable. Hyperpolarized (HP) ^13^C magnetic resonance spectroscopy (MRS) is a promising technique for assessing *in vivo* metabolism. In addition to applications in oncology, the most commonly used probe of [1-^13^C] pyruvate has shown potential in assessing neuroinflammation-linked metabolism in mouse models of multiple sclerosis and traumatic brain injury. Here, we wished to investigate LPS-induced neuroinflammatory changes using HP [1-^13^C] pyruvate and HP ^13^C urea.

2D chemical shift imaging following simultaneous intravenous injection of HP [1-^13^C] pyruvate and HP ^13^C urea was performed at baseline (day 0), day 3 and day 7 post intracranial injection of LPS (n=6) or saline (n=5). Immunofluorescence (IF) analyses were performed for Iba1 (resting and activated microglia/macrophages), GFAP (resting and reactive astrocytes) and CD68 (activated microglia/macrophages).

A significant increase in HP [1-^13^C] lactate production was observed in the injected (ipsilateral) side at 3 and 7 days of the LPS-treated mouse brain, but not in either the contralateral side or saline-injected animals. HP ^13^C lactate/pyruvate ratio, without and with normalization to urea, was also significantly increased in the ipsilateral LPS-injected brain at 7 days compared to baseline. IF analyses showed a significant increase in CD68 and GFAP at 3 days, followed by increased numbers of Iba1 and GFAP positive cells at 7 days post-LPS injection.

In conclusion, we can detect LPS-induced changes in the mouse brain using HP ^13^C MRS, in alignment with increased numbers of microglia/macrophages and astrocytes. This study demonstrates that HP ^13^C spectroscopy holds much potential for providing non-invasive information on neuroinflammation.

## Introduction

Lipopolysaccharide (LPS) is a bacterial endotoxin commonly used to induce neuroinflammation and generate preclinical animal models of the inflammatory response alone^1^, or to help model complex neurological disorders such as Alzheimer’s disease^2^. Upon *in vivo* injection (intravenous^3^, intraperitoneal^4^ or intracranial^5^), LPS causes an inflammatory response, with release of pro-inflammatory cytokines such as tumor necrosis factor alpha, following immune cell recognition of LPS by their cell surface receptors (specifically toll-like receptor 4)^6^. Studies have shown that intracranial injection of LPS in the mouse brain leads to an increased number and activation of microglia^2,7^ as well as increased astrogliosis^8,9^. Further, upon LPS-induced activation, both activated microglia and astrocytes demonstrate metabolic reprogramming, in particular a shift towards increased glycolysis. This has been shown in BV-2 mouse microglia by Voloboueva *et al*.^10^, where LPS induced an increase in lactate production and a decrease mitochondrial oxygen consumption and ATP production, assessed by both biochemical assays and a Seahorse extracellular flux analyzer. Further, Klimaszewska-Lata *et al*.^11^ showed pyruvate dehydrogenase inhibition in the N9 mouse microglial cell line following LPS treatment. Primary astrocytes in culture were similarly activated (by a combinatory LPS/interferon-gamma treatment) by Bal-Price *et al*.^12^ and were shown to produce more lactic acid, as measured by levels in the growth medium. With the knowledge that neuroinflammation is associated with changes in metabolism of immune and glial cells, one could use non-invasive, clinically-translatable metabolic imaging approaches to enable longitudinal monitoring of neuroinflammatory status, as well as response to anti-inflammatory therapies. Magnetic resonance (MR) techniques include ^1^H MR spectroscopy (MRS), and studies using this methodology have suggested that certain ^1^H-visible metabolites, such as myoinositol, choline, and total creatine could be linked to inflammation, for example in multiple sclerosis patients^13^. Increased lactate as detected by ^1^H MRS has also been reported in patients with traumatic brain injury^14^, and preclinical work by Lodygensky *et al*^15^. showed increased lactate in LPS-injected rat pups. ^1^H MRS in general is used widely in the clinic, and hardware and software is already in place for fairly straightforward implementation of novel applications. However, ^1^H MRS is not yet standard clinical procedure for assessment of neuroinflammation, and optimization work for this particular application is ongoing^16^.

^13^C MRS holds great promise for assessing *in vivo* metabolism. It allows identification of steady-state metabolism in the brain after intravenous administration of ^13^C-labeled substrates over >1h^17^, and has been applied in the healthy mouse^18^, rat^19^ and human brain^20^, the healthy aging brain^21^ and AD^22^. However, these methods require long infusions of substrate and extended scan times, which limits clinical translation. Hyperpolarized (HP) ^13^C MRS^23^ is a rapidly expanding alternative imaging technology for visualizing *in vivo* metabolism^24,25^. Acquisition occurs over a matter of minutes, thanks to the 10,000-fold increase in sensitivity over thermal ^13^C MRS. Thus far, HP ^13^C MRS has been particularly informative on the conversion of HP [1-^13^C] pyruvate to [1-^13^C] lactate via the enzyme lactate dehydrogenase, and there has been a focus on applications in cancer^26,27^ and cardiovascular disease^28^. Alongside analysis of enzymatic fluxes, HP probes can be used to assess perfusion. For this purpose, metabolically inactive probes such as HP ^13^C urea can be injected, as demonstrated in tumor imaging by von Morze *et al*.^29^. Further, if co-polarized and administered with pyruvate, HP ^13^C urea can provide a simultaneous readout of metabolism and perfusion, as shown by Lau *et al*.^30^ in the rat heart.

Recently, HP ^13^C MRS has been demonstrated to be applicable to the detection of neuroinflammation in models of multiple sclerosis (MS)^31^ and traumatic brain injury (TBI)^32,33^, where studies have taken advantage of the inflammation-related increase in production of HP lactate. In a mouse model of MS, HP [1-^13^C] lactate was increased in the corpus callosum, and was associated with a significant increase in activated microglia in that region. This increase was then no longer observed in transgenic mice with a deficiency in their ability to activate microglia^31^. HP ^13^C MRS has been applied to both a rat^33^ and mouse^32^ model of TBI. Both studies demonstrated an increase in HP ^13^C lactate/pyruvate ratio following injury, and in mice, the depletion of microglia prevented this increase from occurring.

In this study, we hypothesized that HP ^13^C MRS could be used to visualize the effect of intracranially-administered LPS in the *in vivo* mouse brain. Given the reports that LPS increases glycolysis in brain microglia and astrocytes, we hypothesized that the induced increase in lactate production could be measured by MRS following administration of HP [1-^13^C] pyruvate. We also co-administered HP ^13^C urea in order to control for any changes in perfusion that occurred as a result of the injection, and carried out our study at the clinically-relevant field strength of 3 Tesla. Our results showed that, following intracranial injection of LPS in the mouse brain, HP [1-^13^C] lactate levels and corresponding HP ^13^C lactate/pyruvate ratios were significantly increased at 3 and 7 days post-injection in the ipsilateral (injected) voxel. Moreover, upon normalization to contralateral side, to HP ^13^C urea, or both, increased HP [1-^13^C] lactate levels and increased HP ^13^C lactate/pyruvate ratios were systematically observed at the 7 day timepoint in the ipsilateral side of LPS-injected animals. Importantly, the 7 day timepoint corresponded to the maximum levels of Iba1 (resting and activated microglia/macrophages), and glial fibrillary acidic protein GFAP (resting and reactive astrocytes), as detected by immunostaining. Overall, our results show that ^13^C MRS of HP [1-^13^C] pyruvate and ^13^C urea can successfully visualize the effect of LPS on the mouse brain.

## Materials and Methods

### Animals

All animal procedures were approved by the Institutional Animal Care and Use Committee of the University of California, San Francisco.

A total of 26 mice were used in this study. Eleven mice (male C57BL/6J, 10-12 weeks old, Jackson Laboratories) underwent the imaging protocol described below prior to surgery (Baseline group). These animals were then injected intracranially with either LPS (n=6) or saline (n=5), and the imaging protocol was repeated at 3 and 7 days post-surgery. Separate groups of animals were euthanized at the experimental timepoints (n=5 baseline, n=6 at 3 days, n=6 at 7 days) for histological tissue analysis.

### Intracranial injections

Mice were anesthetized with isoflurane in oxygen (isoflurane: 3% induction, 2% maintenance), and fur shaved from the top of the head. After placement in a stereotactic frame (Stoelting, IL, USA) and on a heating pad, a small incision was made in the skin to expose the skull. Following orientation using the bregma and lambda to ensure the head was flat, a small hole was made in the skull (1.55mm to right and 1mm front of the bregma, depth of 2.8mm), and 5μl of either saline or 1μg/ml LPS in saline (0.005mg; from *E.coli* O111:B4, Sigma Aldrich) injected automatically into the striatum using a Hamilton syringe (**Figure 1A**). The syringe was then removed and the wound sutured closed. On removal from the anesthesia, animals were ambulatory after 5-10 minutes.

**Figure 1.**
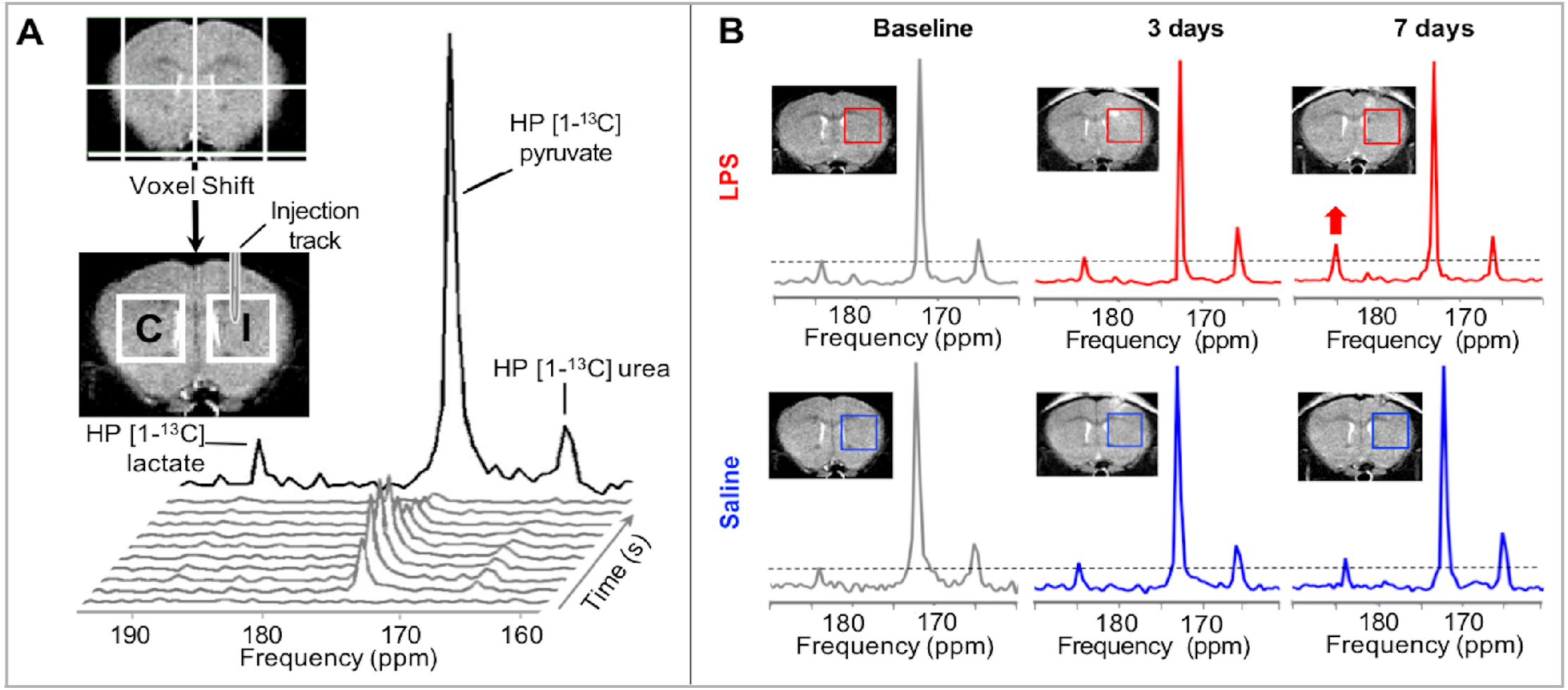
**(A)** MRS grid used for acquisition of HP ^13^C MRSI data overlaid on a T_2_-weighted MR image, and voxels of interest post-voxel shift used for analysis (C=contralateral; I=ipsilateral). Stack of dynamic HP ^13^C spectra (grey), with summed spectrum at the rear (black), showing the resonances of HP [1-^13^C] pyruvate, HP [1-^13^C] lactate and HP ^13^C urea. **(B)** T_2_-weighted MR images and summed HP ^13^C spectra from ipsilateral voxel at each timepoint, for both LPS (red) and saline (blue) animals.

### MR acquisitions

MR acquisitions were performed on a 3 Tesla horizontal MR system (Bruker Biospec) with a dual-tuned ^1^H/^13^C mouse head volume coil (2cm diameter, Doty Scientific, South Carolina). Animals were anesthetized with isoflurane (2% in O_2_) and a 27G tail vein catheter placed before the animal was positioned in a home-built cradle inside the MR system. T_2_-weighted images of the brain were acquired for co-registration of the spectral voxels (field of view (FOV)=20×20mm, matrix 192×192, number of averages (NA)=4, repetition time (TR)=2s, echo time (TE)=60ms, 9 slices of thickness 1mm; total acquisition time 10min 8s). 24μl [1-^13^C] pyruvate sample (pyruvic acid, 15mM OX63 trityl radical (Oxford Instruments), and 1.5mM Gad-DOTA) and 55μl ^13^C urea (6.4M urea in glycerol, with 23mM OX63 trityl radical) were co-polarized for ~1 hour in a Hypersense polarizer (Oxford Instruments, 3.35T), and rapidly dissolved in 4.5ml heated buffer (80mM NaOH in Tris with EDTA) to give a pH7 solution of 80mM pyruvate and 78mM urea. Immediately following dissolution, 300μl of this HP solution was injected via the tail vein catheter over 14 seconds. Data were acquired from the start of intravenous injection every 4.2s using a dynamic 2D chemical shift imaging (CSI) sequence (slice thickness 5mm, FOV=24×24mm, matrix 8×8, TR=66.4ms, TE=1.24ms; total acquisition time 1min 25s).

### Histological analysis

Animals were euthanized using an overdose of ketamine/xylazine, and then perfused with ice-cold phosphate-buffered saline solution (0.9%), followed by an ice-cold paraformaldehyde (PFA, 4%) solution. The brains were removed and submerged in PFA for 2 hours, before being transferred to a sucrose gradient (5% for 2 hours, 10% for 2 hours, 20% overnight). Brains were then frozen in liquid nitrogen and stored at −80°C. Cryosections (10μm) from the imaging voxels were obtained using a microtome (Leica Biosystems, Germany). Immunofluorescence staining was carried out using the following antibodies: a primary rabbit anti-GFAP (resting and reactive astrocytes, 1:500 dilution, Z0334, Dako), and a primary rabbit anti-Iba1 (resting and activated microglia/macrophages, 1:500 dilution, 019-19741, Wako), both with secondary antibodies goat anti-rabbit fluorescein isothiocyanate (1:500 dilution, 111-096-144, Jackson Immunoresearch Lab), and a primary rat anti-mouse CD68 antibody (activated microglia/macrophages, 1:200 dilution, MCA1957, Biorad) with secondary goat anti-rat Alexa-Fluor 555 (1:200 dilution, A21434, Invitrogen). Slides were counterstained using Hoechst 33342 (H3570, 1:2000 dilution; Invitrogen), then sections were mounted using Prolong Gold Antifade (P36930; Invitrogen).

Fluorescence image acquisition was performed using an inverted microscope (Ti, Nikon). The images were recorded with an Andor Zyla 5.5 sCMOS camera at 20x magnification. Quantitative analyses of immunofluorescence images from the striatum posterior to the injection site within the MRS imaging voxel were performed using NIH ImageJ (v2.0.0). Quantification was executed based on coverage, and expressed as percentage of the total area.

### Analysis of hyperpolarized ^13^C data

For each animal at each timepoint, HP ^13^C 2D CSI data were analyzed using in-house MATLAB code. First, voxel shifts were applied to the raw data to generate data from ipsilateral (injection site) and contralateral voxels (symmetrical from midline, **Figure 1A**). Dynamic spectra over time (shown in grey) were summed to generate a summed spectrum (shown in black, **Figure 1A**). HP [1-^13^C] pyruvate, HP [1-^13^C] lactate, and HP ^13^C urea levels were estimated using a Lorentzian fit. HP ^13^C lactate/pyruvate and HP ^13^C pyruvate/urea ratios were calculated, and subsequently HP ^13^C lactate/(pyruvate/urea). Finally, all values (individual metabolites and all ratios) were normalized to data from the contralateral side of the brain to eliminate intra-animal variability.

### Statistical analyses

For all CSI data, a repeated measures one-way analysis of variance (ANOVA) with a Tukey multiple comparisons test was used to establish differences within treatment groups over time, for each side of the brain. Histological data were analyzed using two-way ANOVAs with Tukey multiple comparisons tests to establish significant differences between timepoints and sides of the brain. Graphs show standard deviation, and statistical significance was considered when p≤0.05 (denoted *). Further significances are denoted **p≤0.01, ***p≤0.001, and ****p≤0.0001.

## Results

### HP [1-^13^C] lactate levels, HP ^13^C lactate/pyruvate ratio and HP ^13^C lactate/(pyruvate/urea) ratio are increased in the ipsilateral voxel following LPS injection

**Figure 1B** shows summed HP ^13^C spectra at each timepoint of interest (baseline, 3 days and 7 days) for an LPS (red) and a saline-treated (blue) mouse. The resonances of HP ^13^C urea (163ppm), HP [1-^13^C] pyruvate (171ppm) and its metabolic product HP [1-^13^C] lactate (183ppm) were detected at all experimental timepoints, with increased lactate visible at 7 days in the LPS-treated animal.

On full quantification, our results show that HP [1-^13^C] lactate levels were significantly increased in LPS-treated animals in the injected ipsilateral voxel at both 3 days and 7 days when compared to baseline levels (**Figure 2A**; 3 days: 175 ± 33% of baseline levels, p=0.042; 7 days:195 ± 32% of baseline levels, p=0.036). In contrast, no changes in HP [1-^13^C] lactate levels were observed in the contralateral voxel. In the saline-injected group, HP [1-^13^C] lactate levels remained unchanged in both contralateral and ipsilateral voxels throughout the experimental period. When looking at HP ^13^C urea or HP [1-^13^C] pyruvate levels, no significant changes in the levels of either metabolite were detected over time in either saline- or LPS-injected groups in either contralateral or ipsilateral voxels (**Figure 2A**).

**Figure 2:**
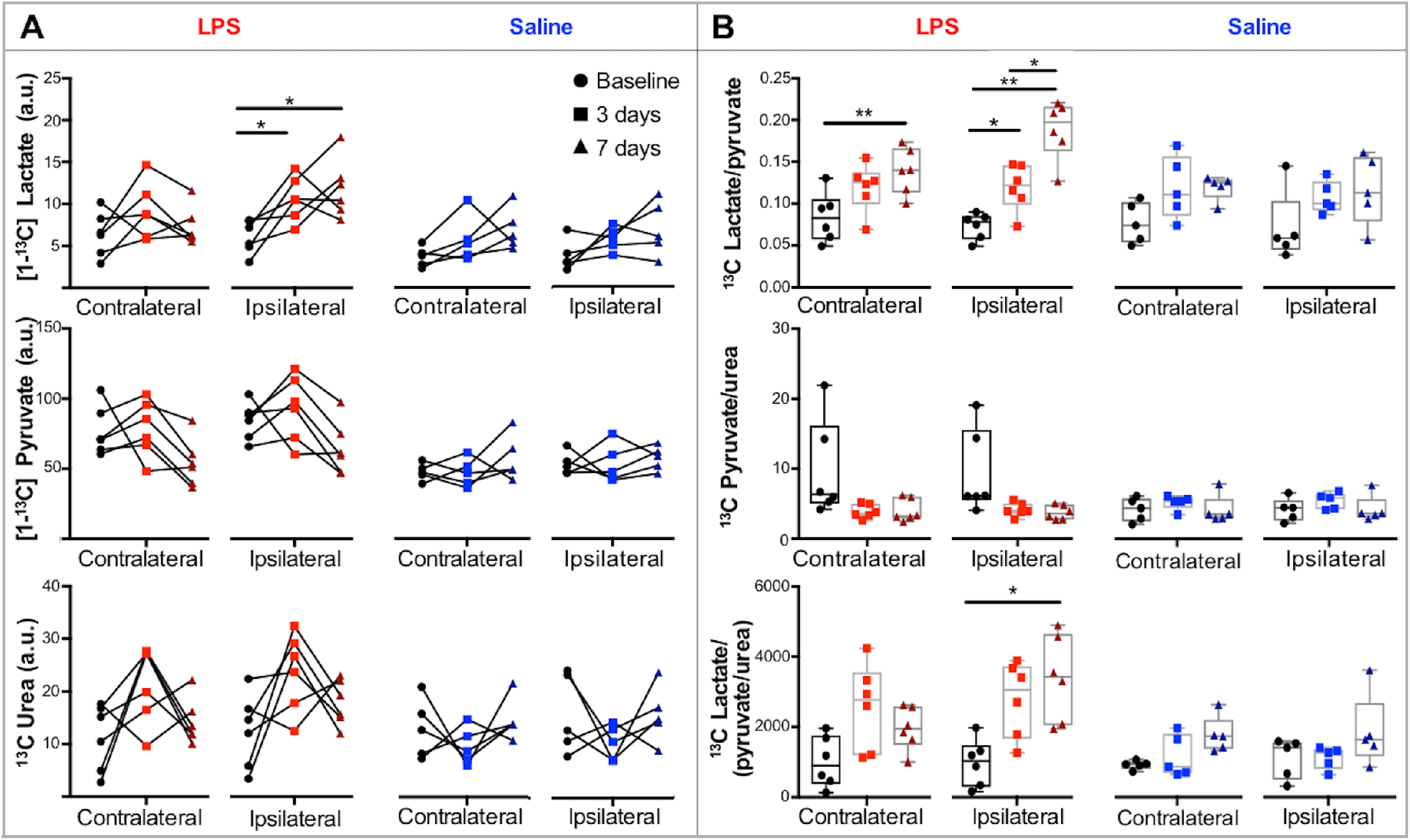
All data are shown for both ipsi- and contra-lateral sides of the brain, in LPS and saline-treated groups, at baseline (•), and at 3 days (■) and 7 days (▲) after surgery. **(A)** Data over time for individual metabolites assessed following injection of HP [1-^13^C] pyruvate. HP [1-^13^C] lactate levels were significantly increased in LPS-treated animals in the injected ipsilateral voxel at both 3 days and 7 days when compared to baseline levels (3 days: 175 ± 33% of baseline levels, p=0.042; 7 days: 195 ± 32% of baseline levels, p=0.036) **(B)** ^13^C Lactate/pyruvate, ^13^C pyruvate/urea, and ^13^C lactate normalized to ^13^C pyruvate/urea ratios. HP ^13^C lactate/pyruvate ratios were significantly increased in the ipsilateral side of the LPS brains at 3 days and 7 days when compared to baseline (3 days: 165 ± 28% of baseline, p=0.046; 7 days: 263 ± 16% of baseline, p=0.001). This ratio was also significantly increased between 3 days and 7 days (7 days: 159 ± 27% of 3 day data, p=0.043). Further, ^13^C HP lactate/pyruvate ratios were significantly increased in the contralateral side of the LPS brains at 7 days compared to baseline data (167 ± 33% of baseline, p=0.004). HP ^13^C lactate/(pyruvate/urea) ratios were significantly increased at 7 days compared to baseline in the ipsilateral side of the LPS-injected animals (347 ± 30% of baseline, p=0.012). *p<0.05, **p<0.01

As shown in **Figure 2B**, HP ^13^C lactate/pyruvate ratios were significantly increased in the ipsilateral side of the LPS brains at 3 days and 7 days when compared to baseline (3 days: 165 ± 28% of baseline, p=0.046; 7 days: 263 ± 16% of baseline, p=0.001). This ratio was also significantly increased between 3 days and 7 days (**Figure 2B**, 7 days: 159 ± 27% of 3 day data, p=0.043). Further, HP ^13^C lactate/pyruvate ratios were significantly increased in the contralateral side of the LPS brains at 7 days compared to baseline data (**Figure 2B**, 167 ± 33% of baseline, p=0.004). In saline-injected animals, no significant changes in HP ^13^C lactate/pyruvate ratios were detected in either ipsilateral or contralateral voxels at any time points.

No significant changes were observed in HP ^13^C pyruvate/urea ratios either in the LPS or saline-treated animals (**Figure 2B**).

Finally, our results show that the HP ^13^C lactate/(pyruvate/urea) ratios were significantly increased at 7 days compared to baseline in the ipsilateral side of the LPS-injected animals (**Figure 2B**, 347 ± 30% of baseline, p=0.012). No significant changes in this ratio could be observed in either the contralateral side of LPS-injected animals, or the ipsilateral and contralateral voxels in the saline-injected animals.

### HP [1-^13^C] lactate levels, HP ^13^C lactate/pyruvate ratio and HP ^13^C lactate/(pyruvate/urea) ratio are increased following LPS injection, upon normalization to contralateral data

Following normalization to contralateral data, our results show that HP [1-^13^C] lactate levels were significantly increased in LPS-treated animals at 7 days when compared to baseline levels and levels at 3 days (**Figure 3A**: 7 days: 164 ± 19% of baseline, p=0.0007; 136 ± 19% of 3 days, p=0.018). In the saline-injected group, HP [1-^13^C] lactate levels remained unchanged throughout the experimental period. When looking at HP ^13^C urea or HP [1-^13^C] pyruvate levels, no significant changes in the levels of either metabolite were detected over time in either saline- or LPS-injected groups (**Figure 3A**).

**Figure 3:**
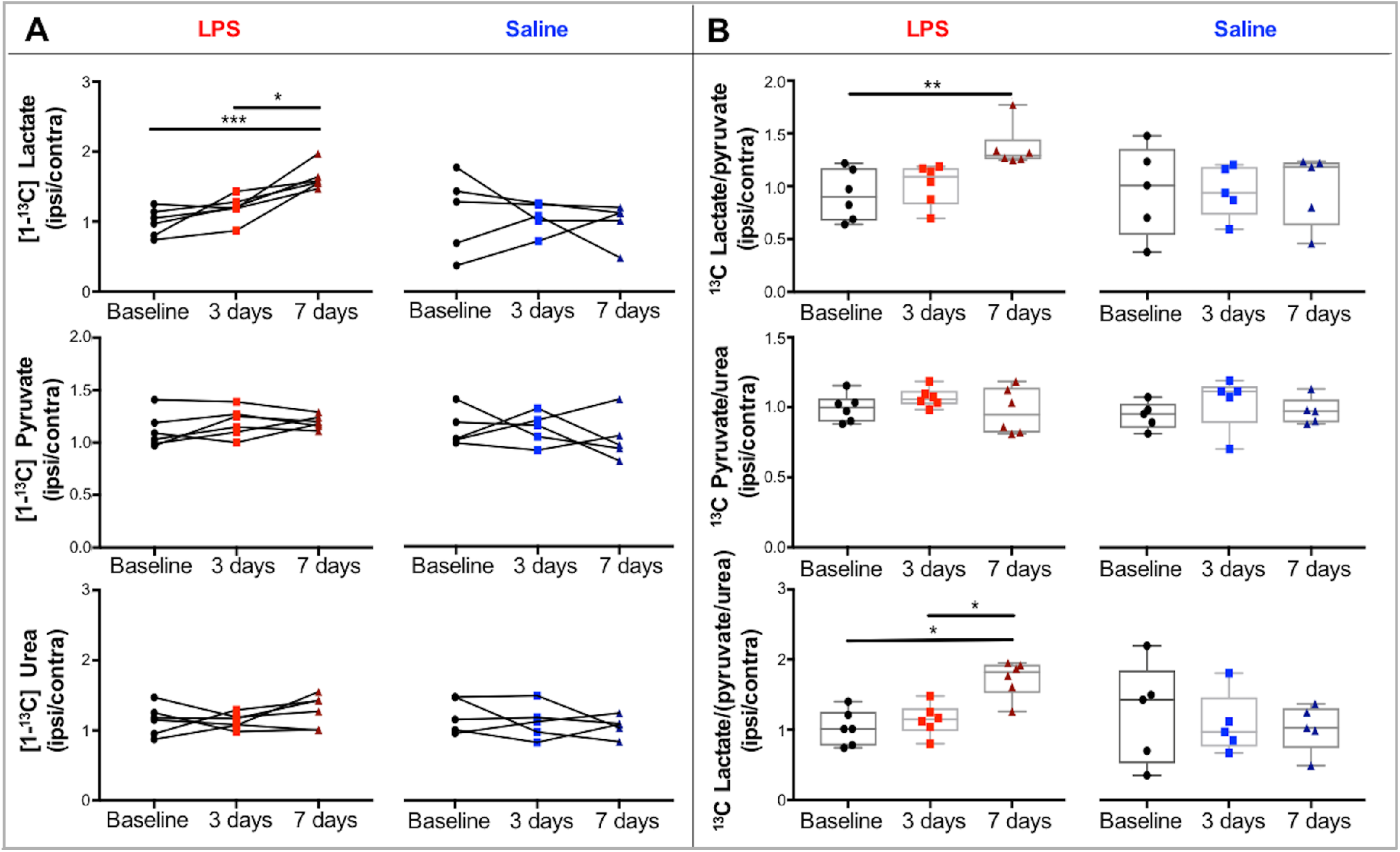
All data are shown for baseline (•), and at 3 days (■) and 7 days (▲) after surgery, in LPS and saline-treated groups. Further, all data were normalized to the contralateral side of the brain. **(A)** Data over time for individual metabolites assessed following injection of hyperpolarized [1-^13^C] pyruvate. HP [1-^13^C] lactate levels were significantly increased in LPS-treated animals at 7 days when compared to baseline levels and 3 day data (7 days: 164 ± 19% of baseline, p=0.0.0007; 136 ± 19% of 3 day data, p=0.018). **(B)** Ratios for ^13^C lactate/pyruvate, ^13^C pyruvate/urea and ^13^C lactate normalized to ^13^C pyruvate/urea. HP ^13^C lactate/pyruvate ratios were significantly increased in the LPS brains at 7 days when compared to baseline (150 ± 28% of baseline, p=0.0097). HP ^13^C lactate/(pyruvate/urea) ratios were significantly increased at 7 days compared to both baseline and 3 day data (7 days: 169 ± 23% of baseline p=0.026, 151 ± 23% of 3 day data, p=0.019). *p<0.05, **p<0.01, ***p<0.001

As shown in **Figure 3B**, HP ^13^C lactate/pyruvate ratios were significantly increased in the LPS brains at 7 days when compared to baseline (7 days: 150 ± 28% of baseline, p=0.0097). In saline-injected animals, no significant changes in HP ^13^C lactate/pyruvate ratios were detected at any time point. **Figure 3B** also shows that we observed no significant changes in HP ^13^C pyruvate/urea ratios in either LPS or saline-treated animals.

Finally, our results show that the HP ^13^C lactate/(pyruvate/urea) ratios were significantly increased at 7 days compared to both baseline and 3 day data (**Figure 3C**, 7 days: 169 ± 23% of baseline p=0.026, 151 ± 23% of 3 day data, p=0.019). No significant changes in this ratio could be observed in the saline-injected animals.

### Iba1, CD68 and GFAP levels are modulated following LPS injection

**Figure 4** shows the results for Iba1 (resting and activated microglia/macrophages), CD68 (activated microglia/macrophages), and GFAP (astrogliosis) immunostaining performed in the ipsilateral (I, large square) and the contralateral side (C, small insert) on LPS-injected animals at baseline, 3 days and 7 days. All values for immunostaining from saline-injected animals were below 0.16% coverage (data not shown).

**Figure 4:**
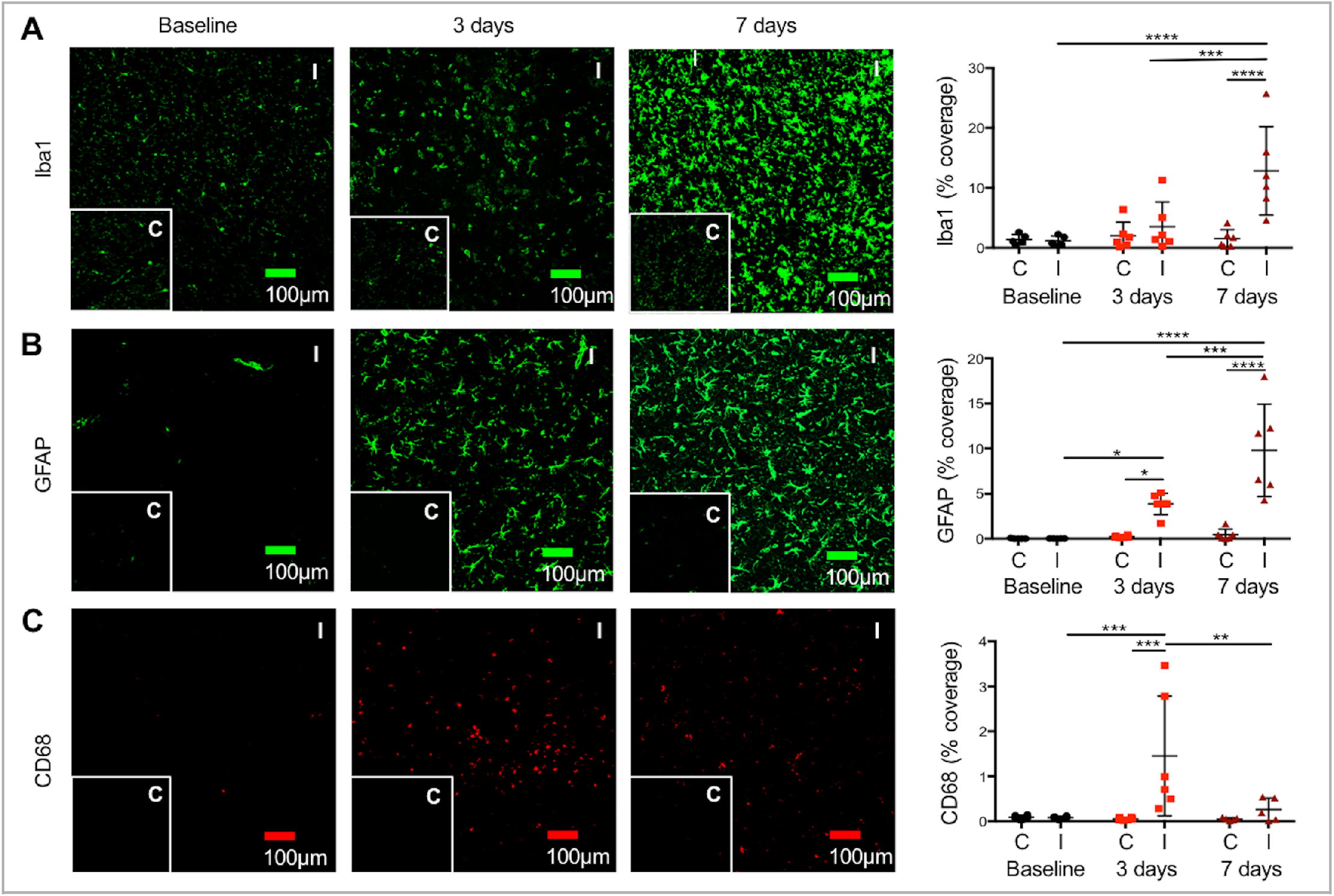
Histological analysis of ipsi- and contra-lateral brain slices at each timepoint in the LPS-treated animals. Panels show Iba1, CD68 and GFAP staining examples (contralateral inset), with quantification of percentage coverage. **(A)** Iba1 staining was significantly increased in the ipsilateral side of LPS-treated animals at 7 days compared to baseline and 3 day levels, and at the 7 day timepoint compared to contralateral (7 days: 1096% of baseline, p<0.0001; 363% of 3 day data, p=0.0006, 835% of contralateral p=0.0001). (B) GFAP staining was significantly increased in the ipsilateral side of LPS-treated animals at 7 days compared to ipsilateral baseline and 3 day levels (7 days: 48550% of baseline, p<0.0001, 254% of 3 days, p=0.0002). Ipsilateral GFAP staining was also significantly higher at 3 days compared to baseline data (3 days: 19130% of baseline, p=0.022 **(C)** CD68 staining was elevated in the ipsilateral side at 3 days post-injection compared to the ipsilateral side at baseline and 7 days (3 day data: 1752% of baseline p=0.0008, 557% of 7 days p=0.003). At 3 days, CD68 was also significantly increased in the ipsilateral side as compared to contralateral (2769% of contralateral, p=0.005). *p<0.05, **p<0.01, ***p<0.001, ****p<0.0001

As shown in **Figure 4A**, Iba1 staining was significantly increased in the ipsilateral side of LPS-treated animals at 7 days compared to baseline and 3 day levels, indicating a large number of microglia/macrophages present at that time point (7 days: 1096% of baseline, p<0.0001; 363% of 3 day data, p=0.0006). At the 7 day timepoint, Iba1 staining was also significantly increased in the ipsilateral side as compared to contralateral (835% of contralateral p=0.0001). No significant differences between ipsilateral and contralateral sides were observed at baseline or 3 days. As depicted in **Figure 4B**, GFAP staining was significantly increased in the ipsilateral side of LPS-treated animals at 7 days compared to ipsilateral baseline and 3 day levels (7 days: 48550% of baseline, p<0.0001, 254% of 3 days, p=0.0002). Ipsilateral GFAP staining was also significantly higher at 3 days as compared to baseline data (3 days: 19130% of baseline, p=0.022). Overall we observed a maximum level of astrogliosis at the 7 day timepoint. When comparing brain sides, our results show that GFAP levels were significantly higher in the ipsilateral voxel than in the contralateral voxel at both 3 days and 7 days (3 days ipsilateral: 1733% of contralateral, p<0.0001; 7 days ipsilateral: 2209% of contralateral, p=0.0003). **Figure 4C** shows that CD68 staining was elevated in the ipsilateral side at 3 days post-injection compared to the ipsilateral side at baseline and 7 days, indicating that monocyte activation reached a maximum level at that time point (3 day data: 1752% of baseline p=0.0008, 557% of 7 days p=0.003). At the 3 day timepoint, CD68 staining was also significantly increased in the ipsilateral side as compared to contralateral (2769% of contralateral, p=0.005). No significant differences between ipsilateral and contralateral sides were observed at baseline or 7 days.

## Discussion

In this study, we demonstrated our ability to image altered brain metabolism following intracranial injection of LPS. Metabolism was assessed using ^13^C MRS following an intravenous injection of HP [1-^13^C] pyruvate and HP ^13^C urea at the clinicially-relevant field strength of 3 Tesla.

Overall, we observed significant changes over time in the detected HP [1-^13^C] lactate, HP ^13^C lactate/pyruvate and HP ^13^C lactate/(pyruvate/urea) only in the LPS-treated group and not in the saline-injected group. This result is indicative of the fact that the metabolic changes observed at 7 days by HP ^13^C MRS were linked to the presence of LPS rather than being a side effect of the surgical procedure.

We did not observe any significant changes in either HP [1-^13^C] pyruvate or HP ^13^C urea at any timepoint at the group level, which indicates that the *in situ* delivery of HP substrates is not significantly different between groups. Recent work by Miller *et al*. has questioned our understanding of cerebral HP signal, and the authors demonstrated a vast increase in metabolic signal when the blood-brain barrier (BBB) is opened by mannitol^34^. The literature discussing BBB leakiness as a result of LPS administration is inconclusive, with a review by Varatharaj *et al*. observing BBB disruption in only 60% of studies considered^35^. Nevertheless, although we have not carried out contrast-enhanced imaging in this study, the fact that we did not observe any significant changes in urea or pyruvate between control and LPS animals suggests that the BBB was not significantly affected by LPS injections^36^.

HP [1-^13^C] lactate alone was significantly increased at 7 days and 3 days compared to baseline in the ipsilateral side of the LPS group. On normalization to HP [1-^13^C] pyruvate, which is typically performed to account for variability in pyruvate injection^26^, the change at 7 days was exacerbated (p=0.001 compared to p=0.036). Not only was ipsilateral HP ^13^C lactate/pyruvate significantly increased at 7 days and 3 days compared to baseline, but further, 7 days was increased compared to 3 days. On normalization to HP ^13^C urea, the 3 day increase in HP ^13^C lactate/pyruvate was no longer significant compared to baseline data. This result may indicate that, although no significant changes were measured when looking at HP ^13^C urea or [1-^13^C] pyruvate/urea data in group analyses, animal specific changes in delivery due to experimental technical variabilities (e.g. injection rate, or percentage polarization of the pyruvate) may have contributed to the increase HP ^13^C lactate/pyruvate ratio observed at the 3 day timepoint. However, at the 7 day timepoint, the observed significant increase in HP [1-^13^C] lactate production remained following normalization, indicating that the increased HP [1-^13^C] pyruvate to HP [1-^13^C] lactate conversion was robust enough to be detected by ^13^C MRS at 3 Tesla.

HP [1-^13^C] pyruvate to HP [1-^13^C] lactate conversion was not expected to be altered in the contralateral side of the brain. We observed no significant changes except for a significantly increased HP ^13^C lactate/pyruvate at 7 days compared to baseline. This unforeseen increase may be due to partial volume effects, as the HP ^13^C lactate/pyruvate ratio is dramatically increased in the adjacent ipsilateral side at that timepoint (263% increase). However, on normalization to HP ^13^C urea, HP ^13^C lactate/(pyruvate/urea) ratio was no longer different between baseline and 7 days, which demonstrates the value of co-injecting HP ^13^C urea to specifically elucidate metabolic changes, independent of experimental variation in intravenous injections.

To provide an internal normalization, and account for biological variability between animals, *in vivo* data were next normalized to the contralateral side of the brain. Once again, significant differences were only observed for HP [1-^13^C] lactate, HP ^13^C lactate/pyruvate and HP ^13^C lactate/(pyruvate/urea) data in the LPS-treated group, not in saline-injected animals, indicating the surgical procedure did not affect brain metabolism. Specifically, upon normalization to contralateral data, differences in HP [1-^13^C] lactate, HP ^13^C lactate/pyruvate and HP ^13^C lactate/(pyruvate/urea) between 7 day and baseline data remained highly significant. Differences in HP [1-^13^C] lactate and HP ^13^C lactate/pyruvate between 3 day and baseline data were no longer observed. For HP [1-^13^C] lactate and HP ^13^C lactate/(pyruvate/urea) parameters, data were significantly different between 3 and 7 days. Overall, data normalized to contralateral brain strengthen the results from the ipsilateral analyses, confirming that HP [1-^13^C] pyruvate to HP [1-^13^C] lactate conversion was increased independently of delivery at 7 days compared to baseline. To confirm the inflammatory response induced by LPS, we carried out histological analysis of animals at each timepoint studied by HP ^13^C MRS. Iba1 and GFAP staining confirmed a substantial increase in microglia/macrophages and astrocytes at 7 days following LPS injection, respectively (compared to both 7 day contralateral data and baseline/3 day ipsilateral data). This result was in line with several previous studies^1,2,7,8,37,38^; Go *et al*^7^ demonstrated a significant increase in microglia at 7 days following an intrahippocampal LPS injection, and Sharma *et al*^38^ showed a similar increase, this time following an intracerebroventricular injection, and additionally observed an increase in GFAP-positive astrocytes.

When considering the histology alongside our *in vivo* data, our results show that the time of maximum HP ^13^C lactate increase as observed by MRS (7 days), coincided with the time of maximal Iba1 and GFAP staining. An increase in number or a change in activation status of these cell types may contribute to the increased HP [1-^13^C] pyruvate to [1-^13^C] lactate conversion. Increased numbers of microglia/macrophages and astrocytes in the ipsilateral LPS-injected brain may be responsible for this increased conversion, given the 1096% and 48550% increase in cell types (respectively), compared to baseline values. Resting microglia express the necessary genes for glycolysis^39^, resting macrophages use both glycolysis and oxidative phosphorylation to produce ATP^40^, and resting astrocytes have been shown to produce lactate in culture^41^. Therefore the increase in number alone may be sufficient to be responsible for the increased MRS signal, irrespective of activation status.

Activation of these glial cells may also play a role. CD68, one of the most commonly-used markers for microglial activation, showed a significant but small increase in the ipsilateral side of the LPS-treated brain at 3 days compared to baseline and 7 days. Interestingly, this timepoint of maximum activation does not coincide with the timepoint of maximum HP [1-^13^C] lactate signal. However, it is recognized that assessing microglial/macrophage activation is not trivial, and likely necessitates measurement of a wide range of markers; such biological characterization is largely beyond the scope of this manuscript. Investigation of the association between CD68 levels and glycolysis would be required to conclude further on the contribution of CD68-activated microglia to the detected *in vivo* HP signal. When considering the contribution of astrocytes, maximum astrogliosis was observed at 7 days (the timepoint of maximum HP [1-^13^C] lactate). Studies have reported that GFAP levels as measured by IF are linked to reactive astrocytes^42–44^. It is thus plausible that astrogliosis as observed at 7 days may contribute to the increased HP ^13^C pyruvate to ^13^C lactate conversion.

Considering activation and cell number, our data suggest that the large increase in number of both cell types (as assessed by percentage coverage in IF) is likely the driving factor for the increased HP ^13^C lactate. However, one cannot distinguish the relative contribution of each cell type to the detected HP [1-^13^C] lactate signal. Studies using modulations of microglia or astrocytes levels could help elucidate cell-specific contributions to the HP signal. Guglielmetti *et al*.^31^ showed increased mononuclear phagocytes in the brain alongside increased HP ^13^C lactate/pyruvate in a multiple sclerosis model, and in a traumatic brain injury model^32^ demonstrated that increased HP ^13^C lactate/pyruvate following injury was no longer observed on microglial depletion. Similarly, Lewis *et al*.^45^ showed increased lactate production in the heart following a myocardial infarction, which was normalized following monocyte/macrophage depletion. Future bioreactor studies^46^ of the individual cell types could also be valuable to decipher the contributions of specific cell types. Only a few HP studies have been carried out on a preclinical, murine-dedicated, cryogen-free 3 Tesla system ^47,48^. This study contributes to the validation of HP acquisitions at this clinically relevant field strength, while taking advantage of the improved gradient strength leading to higher spatial resolution compared to a clinical 3 Tesla. We opted for a 2D CSI dynamic acquisition instead of a single timepoint to remove potential bias from variability in inter-user injection procedures, which might affect the kinetics of HP [1-^13^C] pyruvate and HP [1-^13^C] lactate detected by MRS. In future studies, calculations of rates of conversion (k_pl_) could be carried out, providing sufficient SNR is achieved.

## Conclusion

In conclusion, we have shown that we can successfully detect increased HP [1-^13^C] lactate production *in vivo* following LPS injection into the mouse brain, coinciding with an increased number of microglia/macrophages and astrocytes as visualized by histology. Many diseases have inflammatory components, and HP ^13^C MRS could be a valuable additional tool with which to assess inflammatory status non-invasively and longitudinally.

## Acknowledgements

This work was supported by research grants: NIH R01NS102156, Cal-BRAIN 349087, NMSS research grant RG-1701-26630, Hilton Foundation – Marilyn Hilton Award for Innovation in MS Research #17319, Dana Foundation: The David Mahoney Neuroimaging program, and the NIH Hyperpolarized MRI Technology Resource Center #P41EB013598. Fellowship from the NMSS FG-1507-05297 (to C.G.).

## Abbreviations

LPS: lipopolysaccharide
HP: hyperpolarized
IF: immunofluorescence
Iba 1: ionized calcium binding adaptor molecule 1
GFAP: glial fibrillary acidic protein
CD68: Cluster of Differentiation 68
MS: multiple sclerosis
TBI: traumatic brain injury
NA: number of averages
CSI: chemical shift imaging
PFA: paraformaldehyde

## Bibliography

1. Andersson P-B, Perry VH, Gordon† S. The acute inflammatory response to lipopolysaccharide in cns parenchyma differs from that in other body tissues. Neuroscience. 1992;48(1):169–186.

2. Herber DL, Mercer M, Roth LM, et al. Microglial Activation is Required for Aß Clearance After Intracranial Injection of Lipopolysaccharide in APP Transgenic Mice. J Neuroimmune Pharmacol. 2007;2(2):222–231.

3. Nordgreen J, Munsterhjelm C, Aae F, et al. The effect of lipopolysaccharide (LPS) on inflammatory markers in blood and brain and on behavior in individually-housed pigs. Physiol Behav. 2018;195:98–111.

4. Meneses G, Rosetti M, Espinosa A, et al. Recovery from an acute systemic and central LPS-inflammation challenge is affected by mouse sex and genetic background. Scavone C, ed. PLoS One. 2018;13(8):e0201375.

5. Espinosa-Oliva AM, de Pablos RM, Herrera AJ. Intracranial Injection of LPS in Rat as Animal Model of Neuroinflammation. In: Methods in Molecular Biology (Clifton, N.J.). Vol 1041. Humana Press, Totowa, NJ; 2013:295–305.

6. Fang H, Pengal RA, Cao X, et al. Lipopolysaccharide-induced macrophage inflammatory response is regulated by SHIP. J Immunol. 2004;173(1):360–6.

7. Go M, Kou J, Lim J-E, Yang J, Fukuchi K. Microglial Response to LPS Increases in Wild-Type Mice during Aging but Diminishes in an Alzheimer’s Mouse Model: Implication of TLR4 Signaling in Disease Progression. Vol 479.; 2016.

8. Herber DL, Maloney JL, Roth LM, Freeman MJ, Morgan D, Gordon MN. Diverse microglial responses after intrahippocampal administration of lipopolysaccharide. Glia. 2006;53(4):382–391.

9. Fu HQ, Yang T, Xiao W, et al. Prolonged Neuroinflammation after Lipopolysaccharide Exposure in Aged Rats. Ma D, ed. PLoS One. 2014;9(8):e106331.

10. Voloboueva LA, Emery JF, Sun X, Giffard RG. Inflammatory response of microglial BV-2 cells includes a glycolytic shift and is modulated by mitochondrial glucose-regulated protein 75/mortalin. FEBS Lett. 2013;587(6):756–762.

11. Klimaszewska-ŁAta J, Gul-Hinc S, Bielarczyk H, et al. Differential effects of lipopolysaccharide on energy metabolism in murine microglial N9 and cholinergic SN56 neuronal cells. J Neurochem. 2015.

12. Bal-Price A, Brown GC. Inflammatory Neurodegeneration Mediated by Nitric Oxide from Activated Glia-Inhibiting Neuronal Respiration, Causing Glutamate Release and Excitotoxicity. J Neurosci. 2001;21(17):6480–6491.

13. Chang L, Munsaka SM, Kraft-Terry S, Ernst T. Magnetic Resonance Spectroscopy to Assess NeuroInflammation and Neuropathic Pain. J Neuroimmune Pharmacol. 2013;8(3):576–593.

14. Harris JL, Yeh H-W, Choi I-Y, et al. Altered neurochemical profile after traumatic brain injury: (1)H-MRS biomarkers of pathological mechanisms. J Cereb Blood Flow Metab. 2012;32(12):2122–34.

15. Lodygensky GA, Kunz N, Perroud E, et al. Definition and quantification of acute inflammatory white matter injury in the immature brain by MRI/MRS at high magnetic field. Pediatr Res. 2014;75(3):415–423.

16. Albrecht DS, Granziera C, Hooker JM, Loggia ML. In Vivo Imaging of Human Neuroinflammation. ACS Chem Neurosci. 2016;20:470–483.

17. Bonvento G, Valette J, Flament J, Mochel F, Brouillet E. Imaging and spectroscopic approaches to probe brain energy metabolism dysregulation in neurodegenerative diseases. J Cereb Blood Flow Metab. 2017;37(6):1927–1943.

18. Marin-Valencia I, Hooshyar MA, Pichumani K, Sherry AD, Malloy CR. The ratio of acetate-to-glucose oxidation in astrocytes from a single ^13^ C NMR spectrum of cerebral cortex. J Neurochem. 2015;132(1):99–109.

19. Lanz B, Xin L, Millet P, Gruetter R. *In vivo* quantification of neuro-glial metabolism and glial glutamate concentration using ^1^ H-[^13^ C] MRS at 14.1T. J Neurochem. 2014; 128(1): 125–139.

20. Cheshkov S, Dimitrov IE, Jakkamsetti V, et al. Oxidation of [U-^13^ C]glucose in the human brain at 7T under steady state conditions. Magn Reson Med. 2017;78(6):2065–2071.

21. Boumezbeur F, Mason GF, de Graaf RA, et al. Altered brain mitochondrial metabolism in healthy aging as assessed by in vivo magnetic resonance spectroscopy. J Cereb Blood Flow Metab. 2010;30(1):211–21.

22. Lin AP, Shic F, Enriquez C, Ross BD. Reduced glutamate neurotransmission in patients with Alzheimer’s disease?an in vivo 13C magnetic resonance spectroscopy study. MAGMA Magn Reson Mater Physics, Biol Med. 2003;16(1):29–42.

23. Ardenkjaer-Larsen JH, Fridlund B, Gram A, et al. Increase in signal-to-noise ratio of > 10,000 times in liquid-state NMR. Proc Natl Acad Sci U S A. 2003;100(18):10158–10163.

24. Chaumeil MM, Najac C, Ronen SM. Studies of Metabolism Using (13)C MRS of Hyperpolarized Probes. Methods Enzymol. 2015;561:1–71.

25. Kurhanewicz J, Vigneron DB, Ardenkjaer-Larsen JH, et al. Hyperpolarized 13C MRI: Path to Clinical Translation in Oncology. Neoplasia. 2019;21(1):1–16.

26. Chaumeil MM, Ozawa T, Park I, et al. Hyperpolarized 13C MR spectroscopic imaging can be used to monitor Everolimus treatment in vivo in an orthotopic rodent model of glioblastoma. Neuroimage. 2012;59(1):193–201.

27. Nelson SJ, Kurhanewicz J, Vigneron DB, et al. Metabolic Imaging of Patients with Prostate Cancer Using Hyperpolarized [1-13C]Pyruvate. Sci Transl Med. 2013;5(198):198ra108.

28. Le Page LM, Rider OJ, Lewis AJ, et al. Increasing Pyruvate Dehydrogenase Flux as a Treatment for Diabetic Cardiomyopathy: A Combined 13C Hyperpolarized Magnetic Resonance and Echocardiography Study. Diabetes. 2015;64(8):2735–43.

29. von Morze C, Larson PEZ, Hu S, et al. Imaging of blood flow using hyperpolarized [13C]Urea in preclinical cancer models. J Magn Reson Imaging. 2011;33(3):692–697.

30. Lau AZ, Miller JJ, Robson MD, Tyler DJ. Simultaneous assessment of cardiac metabolism and perfusion using copolarized [1-(13) C]pyruvate and (13) C-urea. Magn Reson Med. January 2016.

31. Guglielmetti C, Najac C, Didonna A, Van der Linden A, Ronen SM, Chaumeil MM. Hyperpolarized ^13^ C MR metabolic imaging can detect neuroinflammation in vivo in a multiple sclerosis murine model. Proc Natl Acad Sci. July 2017:201613345.

32. Guglielmetti C, Chou A, Krukowski K, et al. In vivo metabolic imaging of Traumatic Brain Injury. Sci Rep. 2017;7(1):17525.

33. DeVience SJ, Lu X, Proctor J, et al. Metabolic imaging of energy metabolism in traumatic brain injury using hyperpolarized [1-13C]pyruvate. Sci Rep. 2017;7(1):1907.

34. Miller JJ, Grist JT, Serres S, et al. 13C Pyruvate Transport Across the Blood-Brain Barrier in Preclinical Hyperpolarised MRI. Sci Rep. 2018;8(1):15082.

35. Varatharaj A, Galea I. The blood-brain barrier in systemic inflammation. Brain Behav Immun. 2017;60:1–12.

36. Park I, Lupo JM, Nelson SJ. Correlation of Tumor Perfusion Between Carbon-13 Imaging with Hyperpolarized Pyruvate and Dynamic Susceptibility Contrast MRI in Pre-Clinical Model of Glioblastoma. Mol Imaging Biol. September 2018.

37. Herber DL, Roth LM, Wilson D, et al. Time-dependent reduction in Aß levels after intracranial LPS administration in APP transgenic mice. Exp Neurol. 2004;190(1):245–253.

38. Sharma A, Patro N, Patro IK. Lipopolysaccharide-Induced Apoptosis of Astrocytes: Therapeutic Intervention by Minocycline. Cell Mol Neurobiol. 2016;36(4):577–592.

39. Zhang Y, Chen K, Sloan SA, et al. An RNA-Sequencing Transcriptome and Splicing Database of Glia, Neurons, and Vascular Cells of the Cerebral Cortex. J Neurosci. 2014;34(36):11929–11947.

40. Newsholme P, Gordon S, Newsholme EA. Rates of utilization and fates of glucose, glutamine, pyruvate, fatty acids and ketone bodies by mouse macrophages. Biochem J. 1987;242(3):631–6.

41. Leo GC, Driscoll BF, Shank RP, Kaufman E. Analysis of [1-13C]D-Glucose Metabolism in Cultured Astrocytes and Neurons Using Nuclear Magnetic Resonance Spectroscopy. Dev Neurosci. 1993;15(3–5):282–288.

42. Liddelow SA, Barres BA. Reactive Astrocytes: Production, Function, and Therapeutic Potential. Immunity. 2017;46(6):957–967.

43. Eng LF, Lee YL. Chapter 30: Astrocytic response to injury. Prog Brain Res. 1992;94:353–365.

44. Eng LF, Ghirnikar RS. GFAP and Astrogliosis. Brain Pathol. 1994;4(3):229–237.

45. Lewis AJ, Miller JJ, Lau AZ, et al. Non-Invasive Immuno-Metabolic Cardiac Inflammation Imaging Using Hyperpolarized Magnetic Resonance. Circ Res. February 2018:CIRCRESAHA.117.312535.

46. Radoul M, Najac C, Viswanath P, et al. HDAC inhibition in glioblastoma monitored by hyperpolarized ^13^ C MRSI. NMR Biomed. December 2018:e4044.

47. Qin H, Carroll VN, Sriram R, et al. Imaging glutathione depletion in the rat brain using ascorbate-derived hyperpolarized MR and PET probes. Sci Rep. 2018;8(1):7928.

48. Harris RA, Lone A, Lim H, et al. Cognition and Behavior Aerobic Glycolysis Is Required for Spatial Memory Acquisition But Not Memory Retrieval in Mice. 2019.

